# SELENOT deficiency alters projection neuron migration during corticogenesis in mice

**DOI:** 10.1101/2021.11.23.469805

**Authors:** Emmanuelle Carpentier, Anthony Falluel-Morel, Lisa Brunet, Magalie Bénard, David Godefroy, Loubna Boukhzar, Arnaud Arabo, Youssef Anouar

**Affiliations:** University of Rouen Normandie, INSERM U1239, Neuronal and Neuroendocrine Differenciation and Communication laboratory, IRIB, CURIB, 76821 Mont-Saint-Aignan, France; University of Rouen Normandie, INSERM, the Cell Imaging Platform of Normandy PRIMACEN, IRIB, CURIB, 76821 Mont-Saint-Aignan, France; University of Rouen Normandie, Biological Resource Facility, CURIB, 76821 Mont-Saint-Aignan, France

**Keywords:** corticogenesis, neuronal migration, selenoprotein, antioxidant

## Abstract

During corticogenesis, projection neurons migrate along the radial glial axis to form cortical layers, the alteration of which is associated with functional deficits in adulthood. As byproducts of cell metabolism, reactive oxygen species act as second messengers to contribute to neurodevelopment; however, free radical excess may impede this process. SELENOT is a thioredoxin-like enzyme of the endoplasmic reticulum abundantly expressed during embryogenesis whose gene disruption in the brain leads to neuroblast cell demise due to increased free radical levels. To determine the potential contribution of SELENOT to the establishment of cortical networks, we analyzed first its expression profile in the neocortex at different stages of development. These studies revealed the widespread expression of SELENOT in all cortical layers, and its continous increase throughout mouse lifespan. In addition, we disrupted the SELENOT gene in the cortex using *in utero* electroporation and Nes-Cre/lox knockout. SELENOT deficiency altered neuroblast migration polarity, at the level of radial scaffolding, and projection neuron positionning. These results indicate that SELENOT plays a crucial role during neurodevelopment by sustaining projection neuron migration.

## Introduction

In mammals, the development of the cerebral neocortex is a particular histological process that leads to a laminar six-layered cytoarchitecture. This organization is of a main importance for intracortical and subcortical connectivity in order to ensure the cognitive functions involving this cerebral structure. During corticogenesis between embryonic day (E) 11 and E19 in mice, immature neurons (neuroblasts) are born in the ventricular and subventricular zones (VZ and SVZ) by asymmetric division of radial glial cells (Angevine and Sidman, 1961; Rakic, 1982; Takahashi et al., 1995; Noctor et al, 2001; Noctor et al., 2004). After proliferation, neuroblasts migrate along the radial fibres of radial glial cells from the VZ to the cortical plate (CP) in an “inside-out” manner. Thus, newly generated neuroblasts terminate their migration above oldest neuroblasts to form a new cortical layer (Rakic, 1972; O’Rourke et al., 1992; Nadarajah et al., 2001). The major cell types found in the cerebral cortex are glutamatergic excitatory neurons and GABAergic inhibitory interneurons, that account for approximately 80% and 20% of total neurons, respectively, and this ratio is maintained from development to adulthood (Hendry et al., 1987; Sahara et al., 2012). In contrast to glutamatergic projection neurons which arise from radial glial cells, GABAergic neurons arise from the ganglionic eminence of the telencephalon and migrate tangentially to reach the cortex (Tan et al., 1998; Anderson et al., 1999; Marín et al., 2001). Glutamatergic neuron population is responsible for information treatment and storage through their specific connectivities, depending on the cortical layer where they reside (Kwan et al., 2012), and their activity is regulated by GABAergic neurons. Neuroblast migration abnormalities lead to neurodevelopmental deficits such as autism spectrum disorder (ASD) or schizophrenia (Wegiel et al., 2010; Stachowiak et al., 2017).

Extrinsic and intrinsic factors are involved in proper cell proliferation, migration and differentiation to allow the development of the neocortex. Among these, reactive oxygen species (ROS) which are byproducts of cell metabolism can also act as second messengers to stimulate, for instance, self-renewal of neural progenitors, in specific time-windows and localizations (Le Belle et al., 2011; Chui et al., 2020). However, uncontrolled ROS excess leads to oxidative stress, an aggravating factor of pathologies, which can provoke the loss of the axon initial segment, a decrease in adhesion molecule expression or cell death (Arai et al., 1998; Gogna et al., 2012; Clark et al., 2017).

In order to protect the brain against ROS and to maintain redox homeostasis, numerous antioxidant systems, including key enzymatic activities carried by selenoproteins, are recruited by nerve cells. Selenoproteins are a particular class of enzymes that contain selenium in the organic form of the amino acid selenocysteine (Sec). Selenoproteins are classified in different families, such as glutathione peroxidases (GPX), thioredoxin reductases (TXNRD), and iodothyronine deiodinases (DIO), according to their antioxidant activities mediated by the Sec residue present in their catalytic site. Sec-containing enzymes are crucial for development since their global disruption is lethal in mice (Bösl et al., 1997). Moreover, previous findings showed that selenoproteins are mainly expressed during embryogenesis and that cortical and hippocampal neurons are the favourite site of their expression (Jakupoglu et al., 2005; Zhang et al., 2008). It is now admitted that absence or reduction of selenoprotein expression could contribute to severe neurodevelopmental disorders and neurodegenerative diseases such as ASD, seizure or Parkinson disease (Wirth et al., 2010; Raymond et al., 2014; Cardoso et al., 2015).

The precise functions of certain of these enzymes are not fully elucidated. This is the case of selenoprotein T (SELENOT), a selenoprotein highly conserved during evolution (Moustafa and Antar, 2012), which exerts oxidoreductase activity and whose gene disruption is lethal early during embryonic development in mice (Boukhzar et al., 2016). SELENOT is required for calcium mobilization (Grumolato et al., 2008; Mazza et al., 2019) and plays a role in protein N-glycosylation in neuroendocrine cells (Hamieh et al., 2017). In the brain, SELENOT has neuroprotective effects during development and after injury (Boukhzar et al., 2016; Castex et al., 2016; Alsharif et al., 2021). In the rat neocortex, SELENOT is localised in immature and differentiating neurons (Tanguy et al., 2011) and its conditional gene knockout in the brain causes neuroblast cell death, as well as a decrease in the volume of the neocortex structure. These observations strongly suggest that this selenoenzyme plays an important role during neurodevelopment, in particular for corticogenesis.

To elucidate the role of SELENOT in this developmental process, we sought first to establish its expression pattern in the neocortex to link this expression with cortical development. Then, we genetically disrupted SELENOT expression in cortical neuroblasts harboring a floxed gene through *in utero* electroporation (IUE) of a Cre recombinase construct to evaluate the implication of this important selenoprotein during corticogenesis. The data obtained showed that SELENOT is widely expressed during the different stages of corticogenesis and that its deficiency alters neuroblast migration and projecting neuron positioning.

## RESULTS

### SELENOT is expressed in neocortex throughout the mouse lifespan

In order to understand the contribution of SELENOT to neurodevelopment, we first analyzed its mRNA expression profile in the neocortex from embryonic (E) day 12 to 1-year-old, using laser capture microdissection which allowed isolating different cortical layers. SELENOT mRNA was present at low levels in all cortical layers during embryogenesis, with a higher expression level in the ventricular zone (VZ) at E12 where neuroblast proliferation occurs (Molyneaux et al., 2007). At postnatal stages, SELENOT levels seem to increase compared to embryonic stages although no statistical difference was observed between these stages. We also analyzed SELENOT mRNA levels in older animals, which were significantly higher in all cortical layers at 6 months of age. Indeed, SELENOT mRNA levels exhibited more than 17-fold increase in cortical layers C2-C5 (\**P*<0.05, *n*=6-12) and more than 27-fold increase in cortical layer C6 (\*\**P*<0.01, *n*=6-12) compared to early embryonic (E12 and E18) or postnatal stages (PND0) (Fig. 1A). Using the same samples obtained by laser microdissection, we could reproduce the profile of neuronal nuclei (NeuN or RBFOX3) mRNA abundance, which also displayed an increase during brain maturation (5-6-fold increase for PND90, 6-month-old and 1-year-old compared to younger mice (\**P*<0.05, \**P*<0.05 and \*\**P*<0.01 respectively, *n*=3-12) (Fig. 1B) as previously described (Sekine et al., 2011; Lin et al., 2016).

**Fig. 1:**
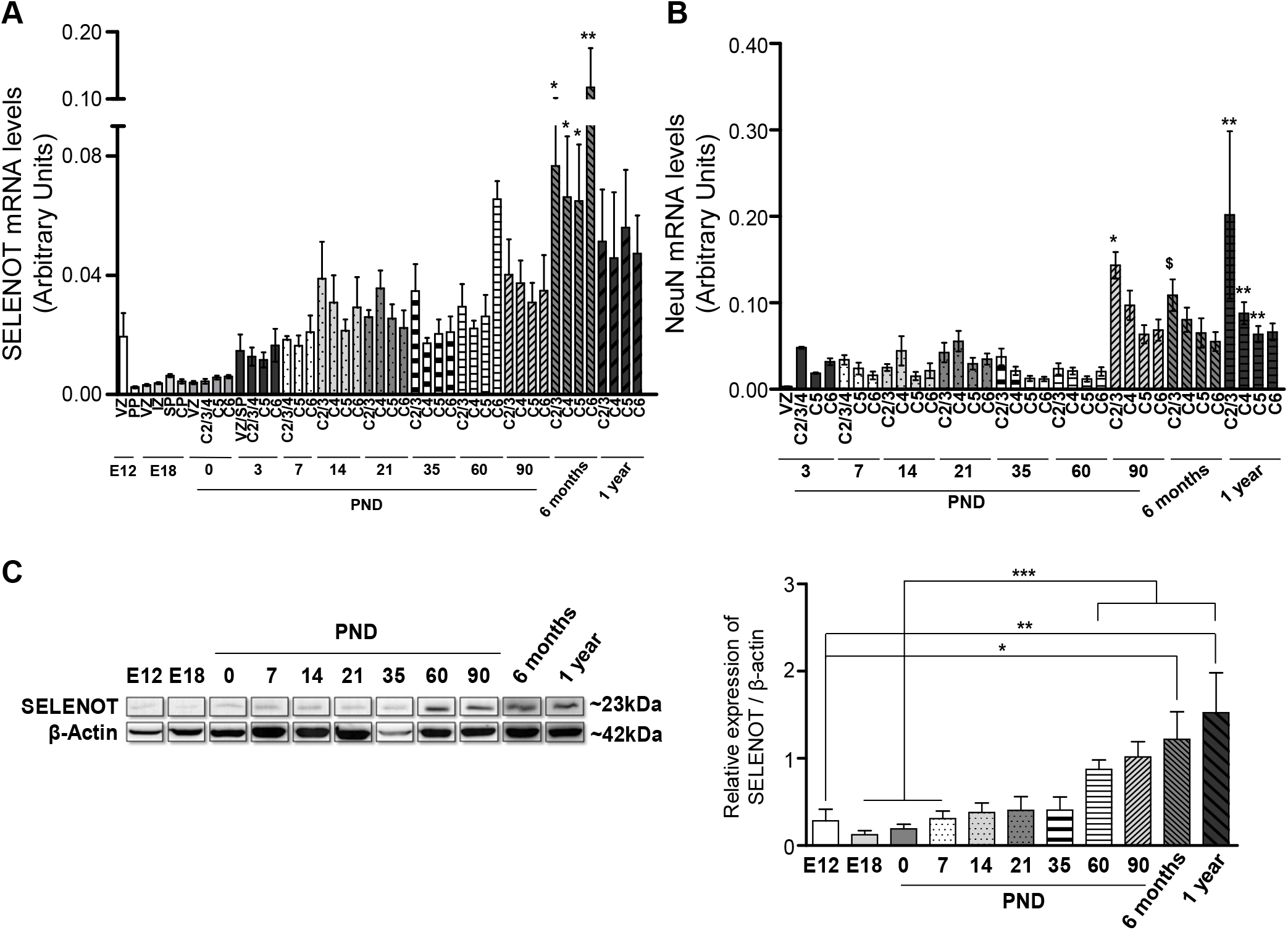
SELENOT is expressed in the neocortex during embryogenesis and throughout the mouse lifespan. (A) SELENOT mRNA levels were analysed in each cortical layer of mice, using laser capture microdissection followed by RT-qPCR, from E12 to 1-year-old. SELENOT mRNA expression was detected in the neocortex during embryogenesis and maturation, with a higher levels in the VZ at E12, becoming significantly higher in aged mice. Relative expression of SELENOT was normalized to GAPDH and represents mean ± sem, *n*=6-12 for E12-1 year. An Increase of >17-fold, \**P*<0.05 for 6 months in C2/3/C4/C5 versus E12 PP, E18 VZ/IZ/SP/CP, PND0 VZ and C2/3/4; >27-fold, \*\**P*<0.01 for 6 months in C6 versus E12 PP, E18 VZ/IZ/SP/CP, PND0 VZ and C2/3/4, using Kruskal-Wallis *U* test and Dunn’s post-test. (B) NeuN relative expression was also analysed in the same samples of microdissected layers, which confirmed previously reported data. Relative expression of NeuN mRNA was normalized to GAPDH mRNA and data were presented as mean ± sem. *n*=3-12 for PND3-1 year. A >6-fold increase, \**P*<0.05 for PND90 C2/3 versus PND3 VZ, PND7 C5, PND14 C2/3/C5/C6 and PND35 C2/3/C4/C5; >5-fold increase, $*P*<0.05 for 6 months C2/3 versus PND14 C5/C6; >5-fold increase, \*\**P*<0.01 for 1-year C2/3/C4/C5 versus PND14 C5/C6 and PND60 C2/3/C4/C5; using Kruskal-Wallis *U* test and Dunn’s post-test. (C) SELENOT protein levels were determined by western blot in the entire mouse cortex, showing its increase in adult and aged mice compared to embryonic and PND stages. SELENOT was normalized to β-Actin and the data represent means ± sem. *n*=3. >3-fold increase, \**P*<0.05 for E12 versus 6 months; >5-fold increase, \*\**P*<0.01 E12 versus 1 year and \*\*\**P*<0.001 for E18/PND0/PND7 versus PND60/PND90/6 months/1 year; using Kruskal-Wallis *U* test and Dunn’s post-test.

SELENOT protein levels were also analyzed to confirm the expression in the neocortex at the different stages. Due to the small size of tissue obtained after laser capture microdissection, we could not analyse the expression of SELENOT in each layer. Therefore, only the entire mouse neocortex could be analysed at different stages. Western blot analysis revealed that SELENOT protein is expressed during embryogenesis at E12 and E18, and increased significantly in adult and aged mice (>3-5 fold increase between PND60, PND90, 6 months and 1 year compared to early embryonic and PND stages (\**P*<0.05; \*\**P*<0.01 and \*\*\**P*<0.001; *n*=3) (Fig. 1C). It should be noted that when SELENOT mRNA and protein expression data were segregated by sex, males and females exhibited similar profiles in all cortical layers and stages analysed (data not shown).

### SELENOT is expressed in proliferating and immature layers within glutamatergic and GABAergic-positive neurons

To better define SELENOT expression profile at the histological level in the neocortex, we used an ultrasensitive *in situ* hybridization analysis. Fig. 2 shows representative SELENOT mRNA staining in all neocortical layers at embryonic stages E14.5 and E17.5, including proliferative (VZ and SVZ), migrating (IZ and SP) and differentiating zones (CP) (Fig. 2A-A’, B-B’). SELENOT mRNA was also expressed in all cortical layers at birth (Fig. 2C-C’), PND 10 (Fig. 2D-D’), PND 20 (Fig. 2, E-E’), PND 60 (Fig. 2F-F’), 6 months (Fig. 2G-G’) and 1 year (Fig. 2H-H’), confirming the PCR results. However, the intensity of SELENOT mRNA signal was similar in brain slices of adult and aged mice compared to embryonic and early postnatal stages (Fig. 2F-F’, G-G’, H-H’ compared to Fig. 2A-A’ to E-E’). In order to determine the neuronal cell types expressing SELENOT gene in the different embryonic cortical layers and to be able to link these data to the following functional studies, we analysed its co-expression with γ-aminobutyric acid (GABA)-ergic and glutamatergic neurons in embryos (E17.5) by *in situ* hybridization or immunohistochemistry.

**Fig. 2:**
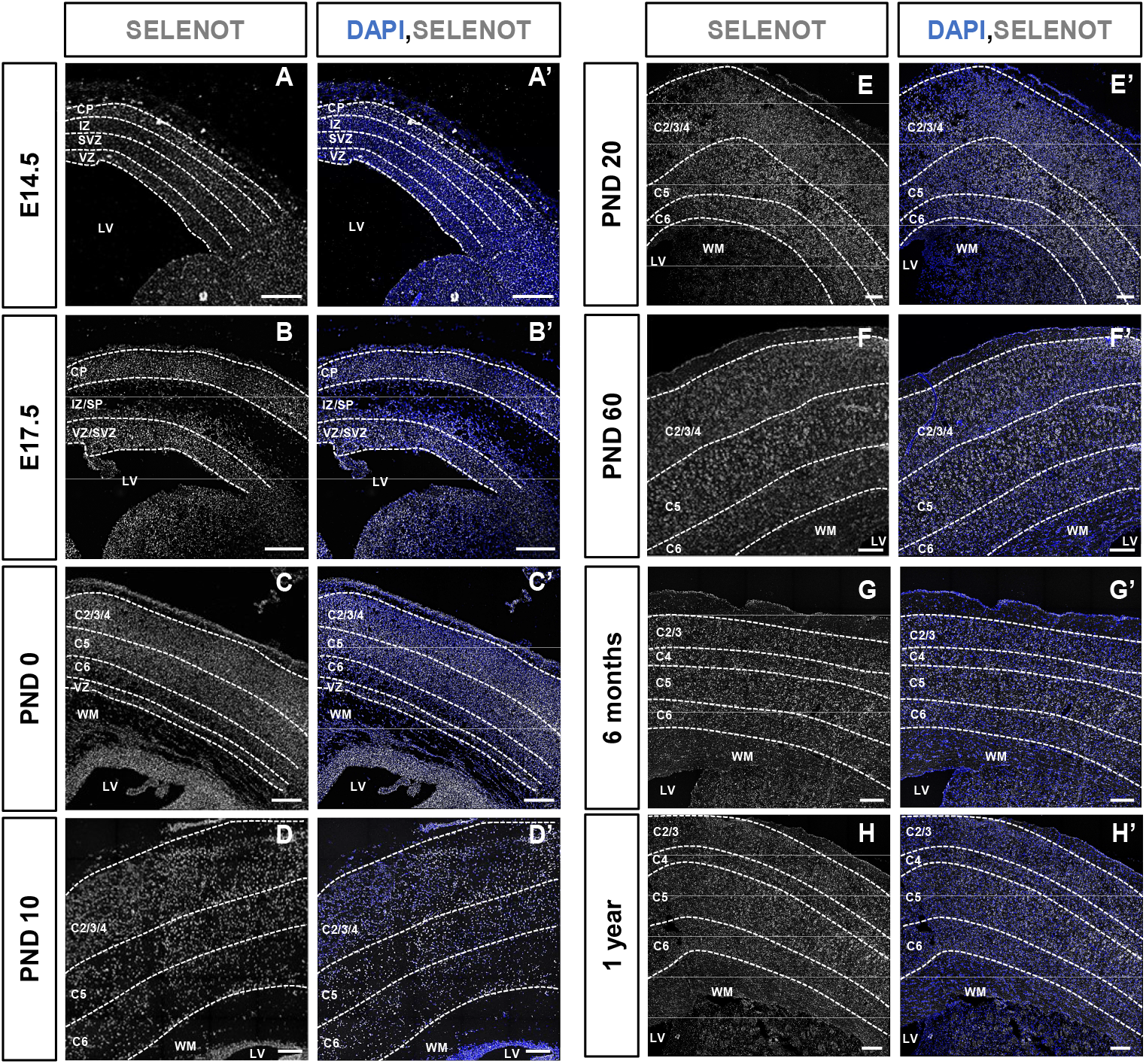
SELENOT mRNA expression in cortical layers throughout mouse development using *in situ* hybridization. Representative images showing SELENOT mRNA from E14.5 to 1-year are presented. SELENOT (grey) is expressed at similar levels in all cortical layers and between embryonic, PND, adult and aged mice (A-H). Staining with DAPI (blue) was used to evaluate cell density (A’-H’). VZ, ventricular zone; SVZ, subventricular zone; IZ, intermediate zone; CP, cortical plate; GE, ganglionic eminence; LV, lateral ventricle; WM, white matter. Scale bar 150 μm.

Co-localization studies showed that SELENOT is present in GABA-positive neurons, labelled with vesicular GABA transporter (VGAT) probes, in the ganglionic eminence where GABAergic neurons are generated, and in the different embryonic cortical layers where GABAergic neuroblasts migrate during corticogenesis (Polleux et al., 2002) (Fig. 3a,b,d). Within cortical layers, SELENOT mRNA signal was present in the VZ where radial glial cells are present and generate neuroblasts (Molyneaux et al., 2007), but also in differentiating neurons labeled with an antibody against doublecortin (DCX), in the CP (Fig. 3a,c,d) (Molyneaux et al., 2007). In addition, SELENOT was present in glutamatergic-positive neurons, stained by type-2 vesicular glutamate transporter (VGLUT2) probes, in all cortical layers (*i.e*. proliferative, migration and differentiation layers) (Fig. 3e,f,g).

**Fig. 3:**
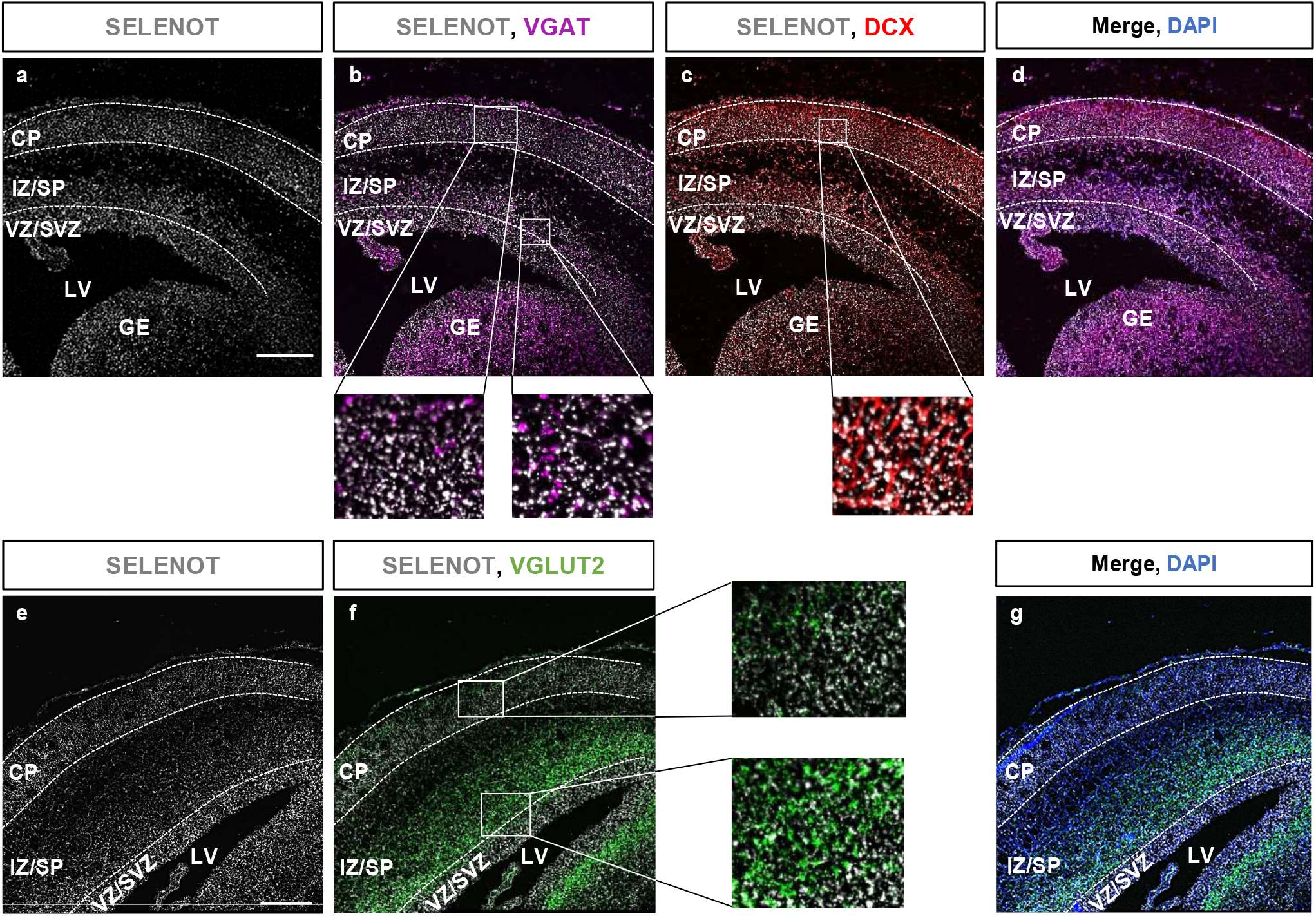
Expression of SELENOT in glutamatergic and GABAergic cells during embryogenesis. Representative images showing the localisation of SELENOT (grey) at E17.5 in the proliferative (VZ/SVZ), the migrating (IZ) and in the CP layers of the mouse neocortex, delimited by dashed lines after DAPI staining (blue) (a,d). SELENOT was detected in GABAergic neurons (VGAT; magenta) in the GE where these neurons are generated and in all cortical layers, where GABAergic neurons migrate (a,b,d). In the cortical plate, SELENOT was found in DCX-positive immature neurons (red), which are differentiating (a,c,d). At E17.5, SELENOT was detected in glutamatergic neurons (VGLUT2, green) in all cortical layers (delimited by DAPI staining, blue) where they proliferate (VZ/SVZ), migrate (IZ) and differentiate (CP) (e,f,g). VZ, ventricular zone; SVZ, subventricular zone; IZ, intermediate zone; CP, cortical plate; GE, ganglionic eminence; LV, lateral ventricle. Scale bars 150 μm.

### Impairment of SELENOT expression in neuronal progenitors through *in utero* electroporation (IUE) leads to neuroblast positioning defects

As SELENOT is expressed during embryogenesis in the neocortex including proliferative and immature layers, we sought to determine its contribution to corticogenesis and more particularly in neuroblast migration. To this intent, we performed IUE in SELENOT^fl/fl^ mice to deliver a plasmid containing a Cre recombinase into the lateral ventricle of embryos at E14.5, when neuroblasts are actively proliferating (Smart and McSherry, 1982; Takahashi et al., 1995). We electroporated the Cre-encoding plasmid in the border of the lateral ventricle in order to disrupt SELENOT gene in neuronal progenitors (Fig. 4A). Both control and pCAG-CRE plasmids carried a gene encoding an enhanced green fluorescent protein (eGFP) to visualize transfected cells (Fig. 4B). Fig. 4C shows the efficiency of the electroporation with green fluorescent cells visible in cortical layers, for the control (Fig. 4Ca,c) and pCAG-CRE (Fig. 4Cd,f) plasmids. The efficiency of SELENOT gene disruption was assayed first by PCR on mice tissue after IUE, and this analysis showed the presence of the SELENOT*^fl/fl^* allele only in the neocortex transfected with the pCAG-CRE plasmid, thus showing the disruption of SELENOT gene (Fig. S1). We also performed immunostaining against SELENOT (Fig. 4Cb,c,e,f) which was detected only in a certain proportion of fluorescent cells (Fig. 4D). Indeed, IUE with the pCAG-CRE plasmid led to a 45% decrease (\*\**P*<0.01; *n*= 522 cells for control and *n*=935 cells for pCAG-CRE condition) in the number of SELENOT-positive fluorescent cells in the neocortex, thus confirming the SELENOT knockout in neuroblasts.

**Fig. 4:**
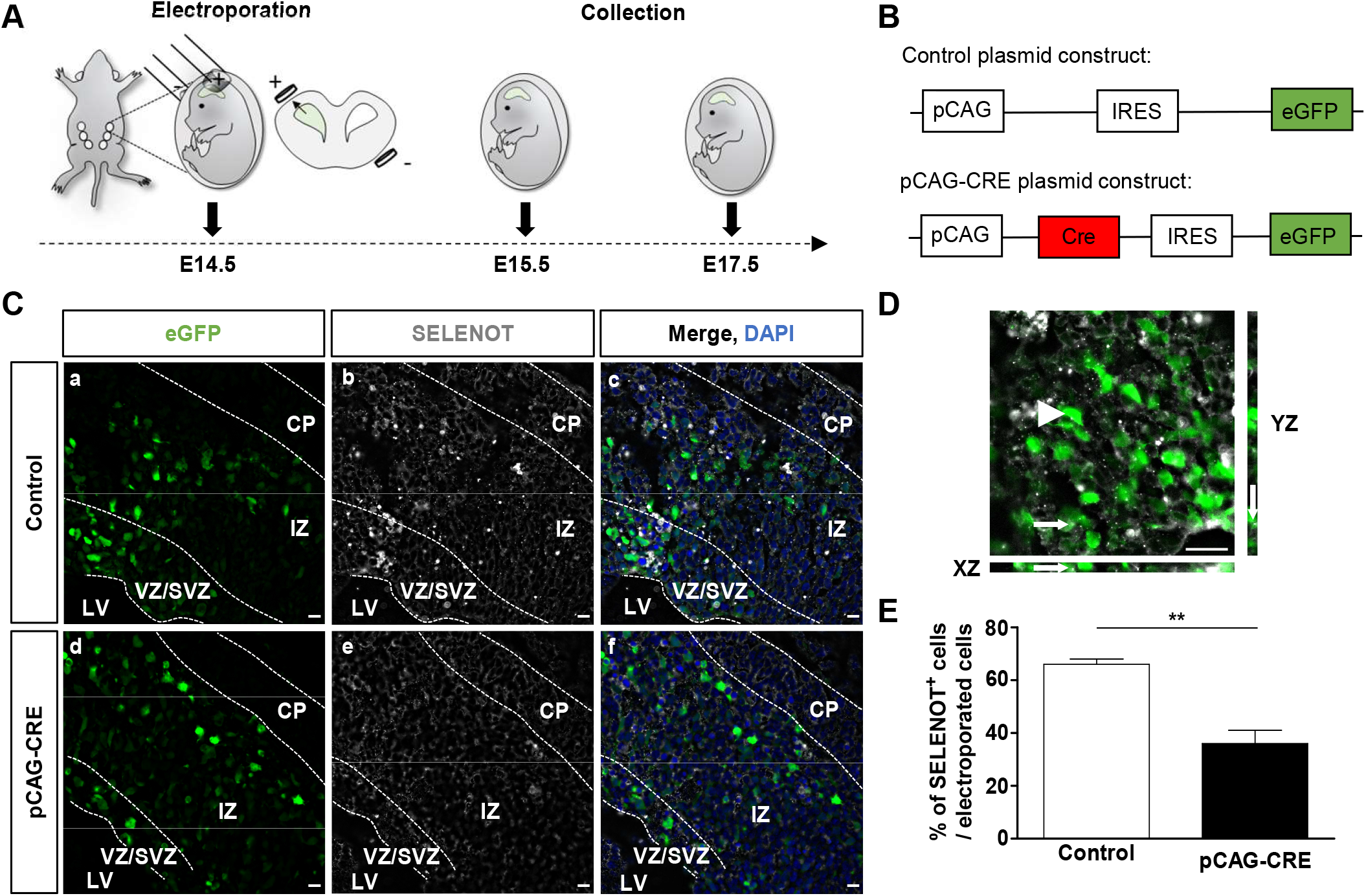
Efficacy of SELENOT disruption by *in utero* electroporation (IUE). (A) IUE was performed in E14.5 embryos where SELENOT gene was floxed (SELENOT*^fl/fl^*). Plasmids were injected in the border of the lateral ventricle of embryo and transfected in neuroblast progenitors by electric pulses. Embryos were collected 24 h (E15.5) or 72 h (E17.5) after IUE. (B) The control plasmid contained a pCAG plasmid backbone with an eGFP sequence to visualise the transfected cells, while the plasmid that disrupts SELENOT gene (pCAG-CRE) contained, in addition, a gene encoding a Cre recombinase which deletes the floxed part of the SELENOT gene in SELENOT*^fl/fl^* embryos. (C) Representative images of transfected cells (green) in E17.5 embryos showing the expression of SELENOT (grey) in all cortical layers (a,b,c) and lower SELENOT labelling in pCAG-CRE-electroporated neocortex (d,e,f). Nuclei were labelled with DAPI. Images in (D) show the colocalization of SELENOT with green-positive cells transfected with the control (arrow) but also its absence in certain transfected green-positive cells (arrow heads) in control condition. Acquisitions were obtained by Z-stack imaging. (E) Quantification of the signal showed 45% decrease of SELENOT-positive fluorescent cells when electroporation was performed with the pCAG-CRE plasmid compared to control. Data are represented as mean ± sem. *n*=522 cells for control and 935 cells for pCAG-CRE conditions, \*\**P*<0.01 using Unpaired *t* test. VZ, ventricular zone; SVZ, subventricular zone; IZ, intermediate zone; CP, cortical plate; LV, lateral ventricle. Scale bars = 10 μm.

Seventy-two hours after IUE (E17.5), most fluorescent cells transfected with pCAG-CRE were observed in the IZ layer, while most fluorescent cells transfected with the control plasmid were located in the CP (Fig. 5A). Quantification of eGFP-positive cells showed a 148% increase (\*\**P*<0.01; *n*=6-7) in the proportion of cells located in the IZ following SELENOT disruption and, conversely, a 37% decrease (\*\**P*<0.01, *n*=6-7) in the percentage of fluorescent cells in the CP (Fig. 5B). Of note, certain neuroblasts expressing pCAG-CRE could reach the CP, probably due to residual expression of SELENOT in these cells.

**Fig. 5:**
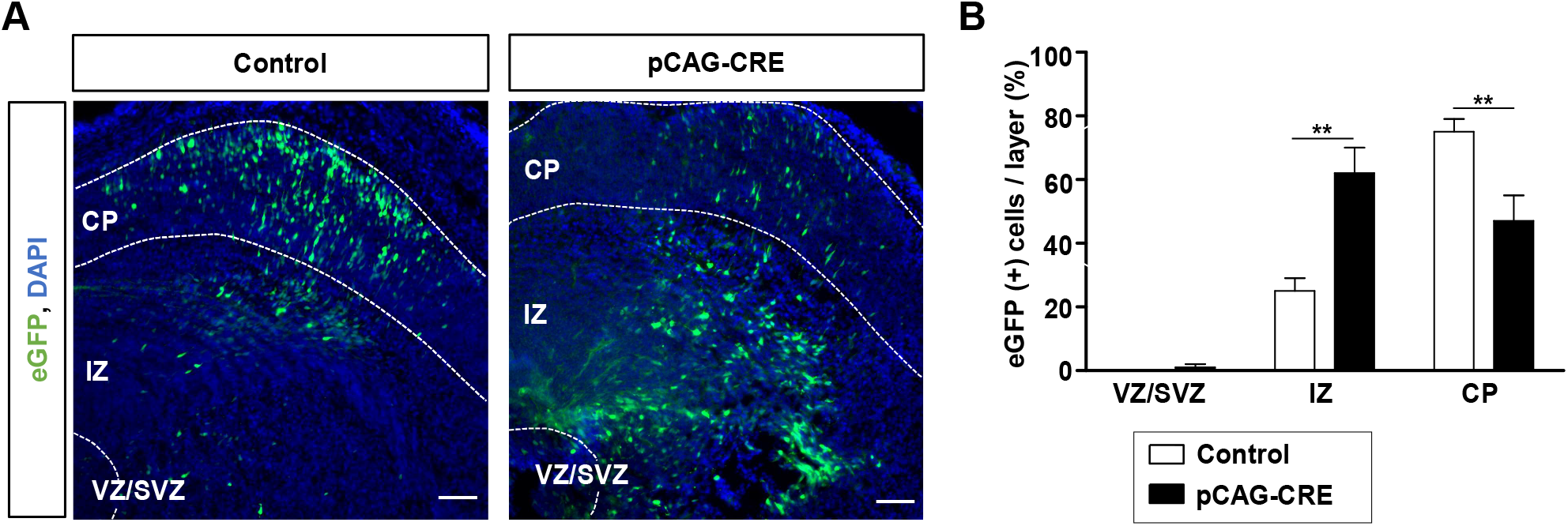
Disruption of SELENOT expression by IUE leads to neuroblast positioning defect. (A) Representative images obtained 72 h after IUE (E17.5) with the control or pCAG-CRE plasmid. In control condition, most eGFP-positive electroporated cells were visible in the CP, while higher proportion of eGFP-positive cells electroporated with the pCAG-CRE plasmid remained in the IZ. Cell nuclei were stained with DAPI (blue). (B) Quantification of eGFP-positive neuroblasts in each cortical layer. Data are represented as mean ± sem. *n*=6 for control and *n*=7 for pCAG-CRE, \*\**P*<0.01 using Mann Whitney *U* test. VZ, ventricular zone; SVZ, subventricular zone; IZ, intermediate zone; CP, cortical plate. Scale bars = 50μm.

### Migrating neuroblast polarity is altered in absence of SELENOT

To better understand the neuroblast positioning defect observed after SELENOT disruption, *i.e*. accumulation of deficient cells in the IZ, we analysed the orientation of their leading process which drives their migration through the radial glia axis up to the neocortex (Nadarajah and Parnavelas, 2002) (Fig. 6A). Measuring the angle (degrees) between the neuroblast leading process and the main axis of radial glial cells (Fig. 6B) and representing the cumulative distribution of these angles in each cortical layer, we found that most SELENOT-deficient neuroblasts display a higher angle of migration in the VZ/SVZ (\**P*<0.05; *n*=12-18 cells per condition) and IZ (\*\*\**P*<0.05, *n*=273-267 cells per condition) compared to cells transfected with the control plasmid (Fig. 6C,D). Neuroblasts that successfully reach the CP have no migration angle alterations (Fig. 6E).

**Fig. 6:**
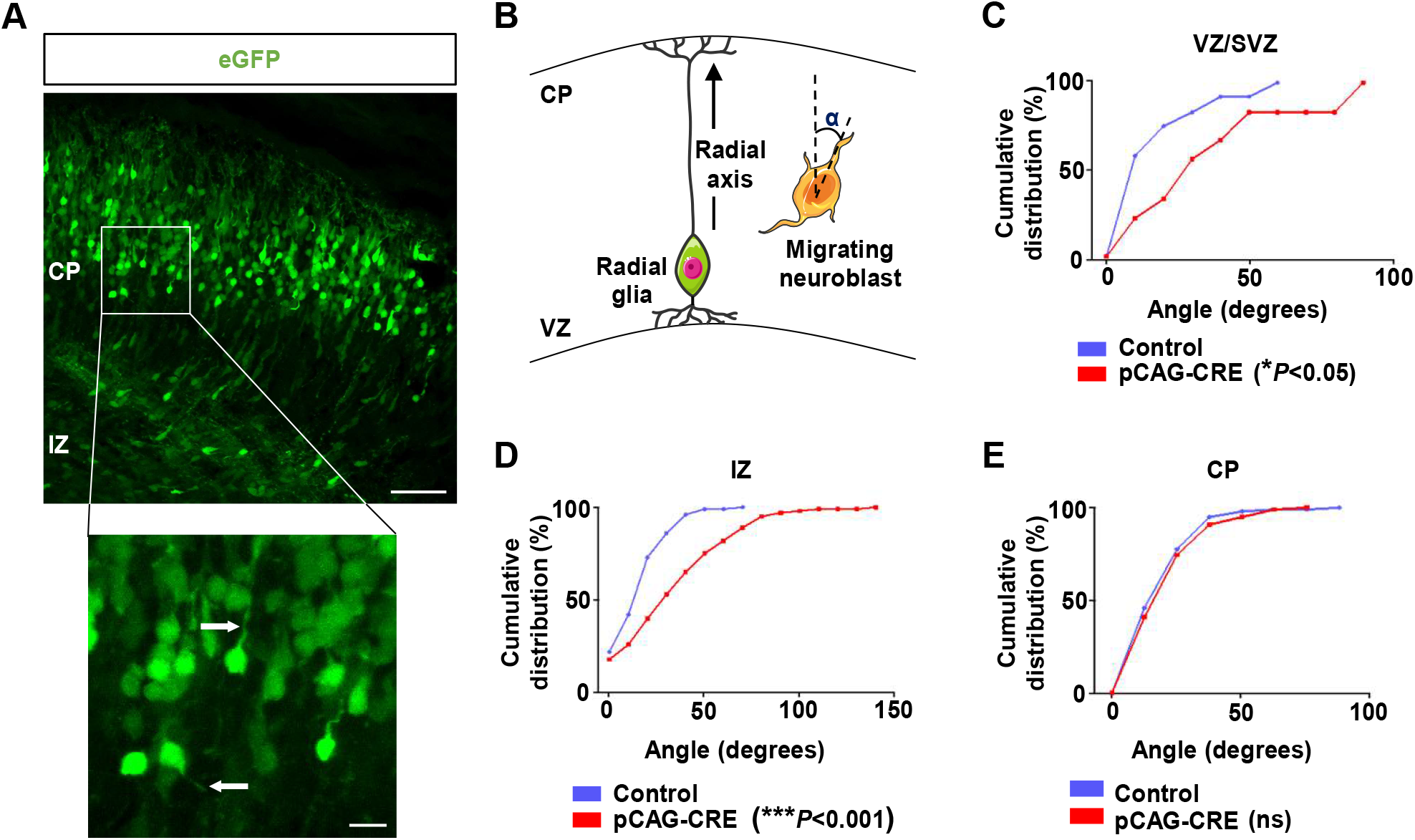
SELENOT deficiency in neuroblasts leads to migration polarity abnormalities. (A) Transfected neuroblasts (green) migrate along the radial glial axis from the VZ to the CP. The inset shows leading processes of neuroblasts (white arrow), which drive their migration. (B) The migration polarity of neuroblasts was determined by calculating the angle (in degrees) between the radial glial axis and the leading process of neuroblasts. The graphs represent the cumulative distribution of the angles measured for cells transfected by the control (blue) or pCAG-CRE (red) plasmids. Seventy-two hours after electroporation, pCAG-CRE cells have a significantly higher angle of migration in VZ/SVZ (C) (*n*=12-18 cells per condition) and IZ (D), (*n*=267-273 cells per condition), compared to control cells. (E) Cells that reach the CP display no difference between the two groups (*n*=117-292 cells per condition). Only cells with a distinguishable leading process were included in this analysis. \**P*<0.05; \*\*\**P*<0.001; ns, not significant versus the respective control using Mann-Whitney *U* test. VZ, ventricular zone; SVZ, subventricular zone; IZ, intermediate zone; CP, cortical plate. Scale bars = 50μm and zoom = 10 μm.

### Absence of SELENOT leads to radial axis abnormalities

Since IUE was performed in the border of the lateral ventricle at E14.5, the plasmid is transfected in radial glial progenitors which serve as a scaffold for neuroblast migration (Rakic, 1972). We then hypothesized that alteration of neuroblast migration could be due to disruption of radial glial scaffolding after SELENOT gene knockout. To investigate this possibility, we first confirmed the localization of SELENOT in radial glia cells in both their soma and processes, using an antibody directed against nestin (Fig. 7A). Then, we used nestin labelling of radial glia cells to analyse the size of their processes in the cortical layers 24 h after IUE of Cre-expressing or control plasmids. Although the full length of each process could not be determined in the tissue sections analysed, we could quantify the apparent length of these processes within each layer of SELENOT-deficient compared to control animals (Fig. 7B). This analysis revealed a decrease of 46% in this parameter in VZ/SVZ (\*\*\**P*<0.001, *n*=8-9 processes), 20% in IZ (\**P*<0.05, *n*=13 processes) and 33% in CP (\*\**P*<0.01, *n*=8-9 processes) (Fig. 7C).

**Fig. 7:**
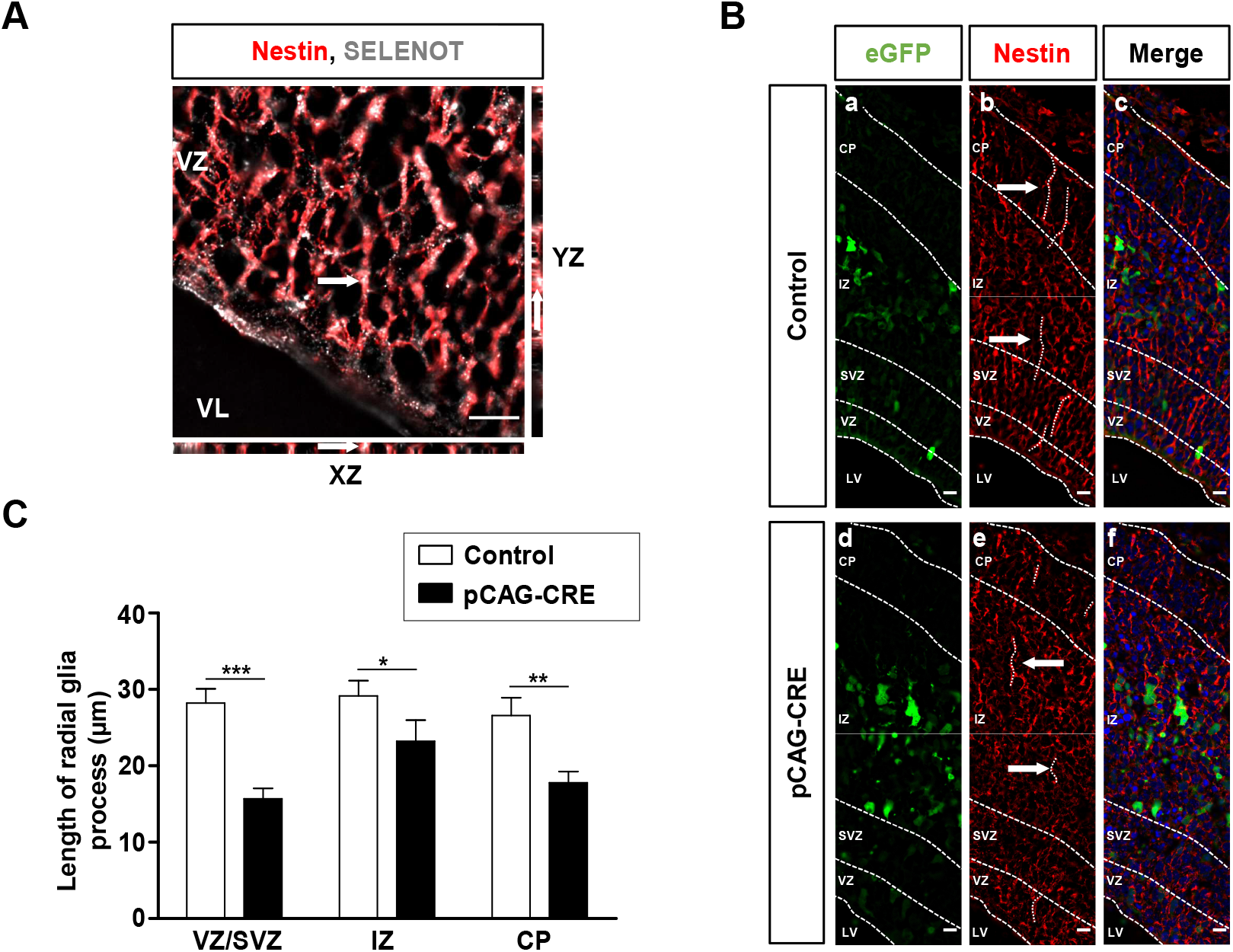
SELENOT disruption in radial glial progenitors leads to alteration of radial glial scaffolding. (A) Immunostaining for nestin (red) was performed in control condition to confirm the expression of SELENOT (grey) in radial glial soma and processes. Z-stack imaging shows the colocalization of SELENOT with nestin (white arrow). (B) Radial glial scaffolds in neocortical slices of E17.5 mice were compared between cells transfected (green) with control (upper panel; a,b,c) or pCAG-CRE (lower panel; d,e,f) plasmids. Radial glial processes have a polarized organization toward the pial surface in the normally developing neocortex (b), but are more disorganised and shorter in pCAG-CRE-treated neocortex (e). The images c and f show the merge of transfected cell fluorescence, nestin labelling and DAPI staining. (C) Quantification of radial glial processes length indicates a decrease of radial glial processes throughout the different cortical layers after transfection with pCAG-CRE plasmid, with 46% decrease in VZ/SVZ (*n*=8-9 processes), 20% decrease in IZ (*n*=13 processes) and 33% decrease in CP (*n*=8-9 processes). Quantification was realized on Z-stack projection and represent means ± sem. \**P*<0.05, \*\**P*<0.01, \*\*\**P*<0.001 using Mann-Whitney *U* test. VZ, ventricular zone; SVZ, subventricular zone; IZ, intermediate zone; CP, cortical plate; LV, lateral ventricle. Scale bars = 10μm.

### SELENOT is required for the proper positioning of deep neurons in adult neocortex

As SELENOT deficiency after IUE alters neuroblast polarity as well as radial glial scaffolding, we sought to determine the positioning of SELENOT-deficient neurons in the neocortex of Nes-Cre/SELENOT*^fl/fl^* adult mice, where SELENOT was disrupted in neural cells. To do this, we immunostained the neocortex with an antibody directed against BAF Chromatin Remodeling Complex Subunit (BCL11B or CTIP2), a transcription factor primarily expressed in neurons of the deep layers (V and VI), and preferentially in layer V (Hevner, 2007; Chen et al., 2008). Indeed, CTIP2 immunolabelling was observed in layers V and VI of the adult neocortex in Nes-Cre/SELENOT^+/+^ mice, with a stronger labelling in layer V (Fig. 8Aa). In contrast, CTIP2-positive cells were more abundant in layer VI and less in layer V in adult SELENOT-deficient mice (Fig. 8Ab). Analysis of CTIP2 distribution in adult Nes-Cre/SELENOT*^fl/fl^* neocortex showed a decrease of 5% and 23% in the C4 and C5 layers (\**P*<0.05; \*\**P*<0.01, *n*=3), respectively, and an increase of 17% in the C6 layer (\*\*\**P*<0.001, *n*=3), compared to adult Nes-Cre/SELENOT^+/+^ mice (Fig. 8B). These data confirm the results obtained with IUE, thus demonstrating that SELENOT plays a physiological role in cortical lamination.

**Fig. 8:**
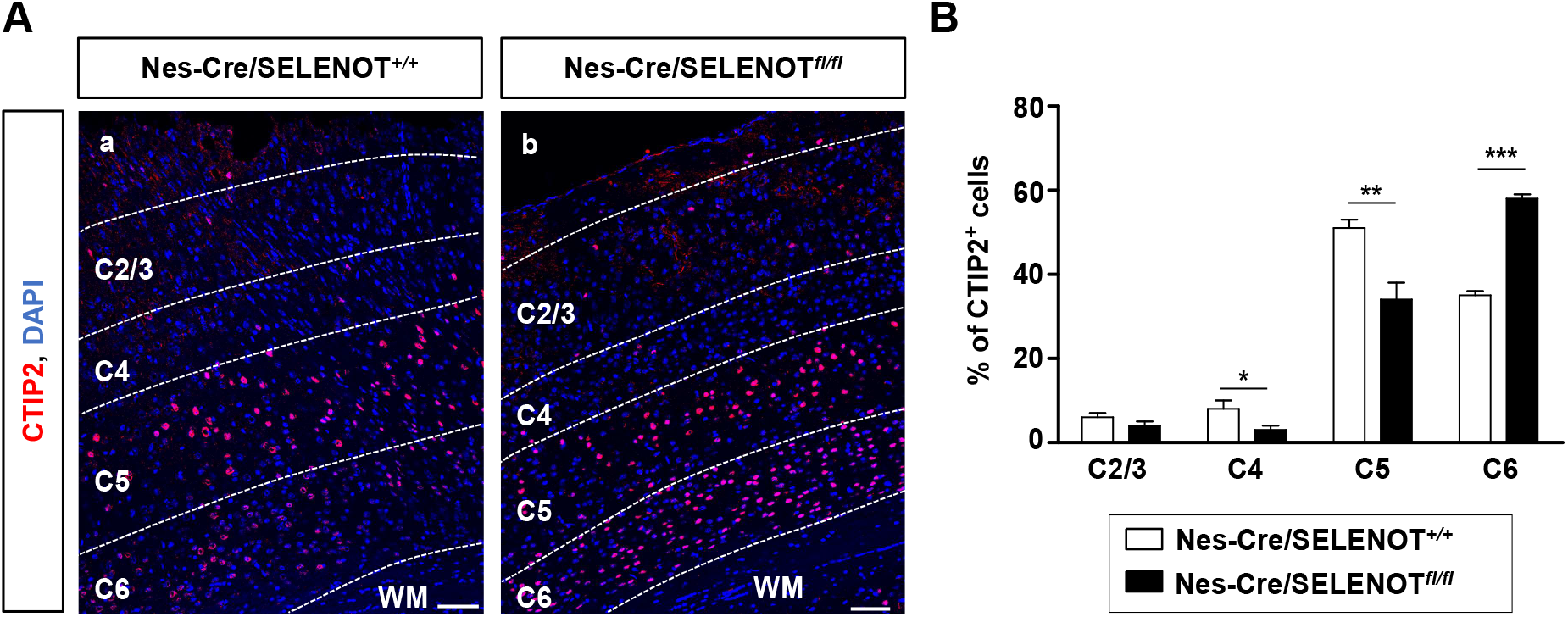
Neurons of deep cortical layers have positioning defects in adult brain SELENOT-deficient mice. (A) Representative images of the neocortex of Cre recombinase-expressing WT (Nes-Cre/SELENOT^+/+^) (a) or Nes-CRE/SELENOT*^fl/fl^* (b) adult mice, labelled for CTIP2, a marker of deep cortical neurons, showing more CTIP2-positive neurons in C6 layer of Nes-Cre/SELENOT*^fl/fl^* mice compared to Nes-Cre/SELENOT^+/+^, and in contrast, less CTIP2-positive neurons in C5 and C4 layers. Cell nuclei are stained with DAPI (blue). (B) Quantification of CTIP2-positive neurons in each cortical layer indicates a 17% increase in C6 and a 5% and 23% decrease in C4 and C5, respectively, in Nes-Cre/SELENOT*^fl/fl^* compared to Nes-Cre/SELENOT^+/+^ neocortex. Data are presented as mean ± sem (*n*=3), \**P*<0.05, \*\**P*<0.01, ***P<0.001 using Unpaired *t* test. C2/3/4/5/6, cortical layer 2/3/4/5/6; WM, white matter. Scale bars = 50μm.

## Discussion

### SELENOT is expressed in the cerebral cortex during mouse lifespan

Previous studies have shown a role of SELENOT in the ontogenesis of the brain in mice (Castex et al., 2016), but its precise function in the different cerebral ontogenetic processes such as corticogenesis remains unknown. Here, we report the importance of this selenoenzyme for proper neuroblast migration during corticogenesis. To understand its contribution for the development of the different cortical layers, we first analysed SELENOT expression profile in mouse cerebral cortex, using biochemical (*i.e*. qPCR following laser capture microdissection, western blot) and histological (*i.e*. RNAscope *in situ* hybridization) methods. Our data indicate that SELENOT is expressed in the mouse brain early during embryogenesis, confirming our previous results obtained in the rat (Tanguy et al., 2011). However, this latter study showned that the global expression of SELENOT in the rat brain declines after birth and in adulthood. In the mouse cerebral cortex, our present data showed an inverse trend with increased expression of SELENOT in postnatal days and in adulthood. Moreover, quantitative PCR data showed that SELENOT mRNA expression is similar in cortical layers at each stage. RNAscope *in situ* hybridization confirmed the presence of SELENOT mRNA in all layers of the mouse neocortex from embryogenesis to 1 year-old.

In addition, SELENOT expression significantly increased in the neocortex of aged mice (6 months and 1-year old). Our previous data showned that SELENOT can protect neuroblasts and neurons against oxidative stress (Boukhzar et al., 2016; Castex et al., 2016; Abid et al., 2019), suggesting that the rise in SELENOT expression observed in cortical layers of aged mice is associated to a defense mechanism against ROS. In fact, the aging brain is exposed to various insults, including ROS which can trigger cell senescence (Perez-Campo et al., 1998; Passos et al., 2010). Of note, this increase could not be visualized by RNAscope *in situ* hybridization. This apparent discrepancy could be due to the tissue architecture, the density of cells or the intracellular levels of SELENOT expression in each layer at different stages. Overall, these gene expression results strongly suggest that SELENOT could play a role in the neocortex during its ontogenesis but also later during brain aging.

### SELENOT is present in proliferative and differentiating neuroblasts

RNAscope *in situ* hybridization analysis showed that SELENOT is present in the ventricular and cortical plate layers. This finding suggests that the selenoprotein could be an important actor in the ventricular zones where cell proliferation occurs during embryogenesis. It has been shown that SELENOT expression is crucial to maintain nerve cell division, by promoting cell cycle G1-S transition (Shao et al., 2019). Moreover, the presence of SELENOT in DCX-labeled immature neurones that reside in the cortical plate at E17.5, may also reflect the importance of this protein for neuroblast differentiation. In accordance, previous studies have shown that SELENOT is overexpressed in differentiating cells and participates to calcium mobilization, a process necessary for cell differentiation and specification (Grumolato et al., 2008; Tonelli et al., 2012).

We found that SELENOT mRNA is present in both glutamatergic and GABAergic neuroblasts. Glutamatergic cells expressed SELENOT in all cortical layers, comprising proliferative (VZ, SVZ), migrating (IZ) and differentiating (CP) layers, at E17.5. In fact, glutamate is essential for different developmental processes, depending on the cell localisation. Thus, glutamate stimulates neuroblast proliferation in the VZ, but not in the SVZ (LoTurco et al., 1995; Haydar et al., 2000; Luk and Sadikot, 2004), and neuroblast migration toward the CP, via a chemoattractant effect mediated by NMDA receptors and increased intracellular calcium (Behar et al., 1999; Simonian and Herbison, 2001). However, another study showed that the stimulation of NMDA receptors in the cortex acts as a stop signal for migration (Kihara et al., 2002). These results indicate that glutamate exerts opposite effects during corticogenesis, depending on the time-window and neuroblast localisation. SELENOT ought to play a role in glutamatergic cells during corticogenesis probably to allow appropriate production of this essential neurotransmitter. SELENOT was also detected in GABAergic neurons, which are generated in ganglionic eminences in the ventral telencephalon and which reach the cortex by tangential migration (Polleux et al., 2002). SELENOT is detected in the site of origin of these cells and in generated and migrating GABAergic neuroblasts, at E17.5. These data support a role of SELENOT also in the generation and implementation of GABAergic neurons. Expression of SELENOT in neuronal cells in the developing cortical layers could be involved in the protection of different cortical cells against oxidative stress which is inevitably produced during the different processes, *i.e*. cell proliferation, migration and differentiation, of cortical lamination.

### SELENOT is required for neuroblast migration

To elucidate the possible contribution of SELENOT during corticogenesis, we performed *in utero* electroporation of a Cre-expressing plasmid in mice to inactivate SELENOT gene in floxed neuroblast progenitors. We found that significantly less neuroblasts reach the CP after SELENOT inactivation compared to control, and numerous SELENOT-deficient cells remained in the IZ. Nevertheless, not all transfected cells were impacted since a certain number could be observed at the superficial layer. This could be ascribed to the level or timing of Cre recombinase expression in transfected cells, as attested by different eGFP staining levels among the cells.

During corticogenesis, radial glial cells give rise to neuroblasts and extend long processes known as the radial axis, which serve as a scaffolding for neuroblast migration until the CP (Rakic, 1972). Neuroblasts are guided by the radial axis, notably thanks to adhesive molecules (Anton et al., 1999; Sanada et al., 2004). Since SELENOT-deficient neuroblasts exhibited an altered positioning during corticogenesis, we investigated the angle of neuroblast migration with respect to the radial axis. We found that these neuroblasts located in the VZ/SVZ and IZ have a higher angle of migration compared to control. This means that neuroblasts do not migrate properly towards the CP, and can explain the increase in the number of cells accumulating in the IZ. This abnormality could result from a loss of adhesion between neuroblasts and the radial axis or even from glial scaffolding alterations. In fact, SELENOT is expressed in radial glial cells and its disruption could affect scaffolding. Therefore, we analyzed glial processes morphology using nestin as a marker, and found a decrease in their apparent length with a disorganised scaffold. It can thus be concluded that neuroblast migration is impaired due to a loss of neuroblast polarity, probably as a result of radial glial scaffolding impairment. Interestingly, a role of SELENOT in cell adhesion has been previously described in fibroblasts (Sengupta et al., 2009), and may also occur in neuroblasts and account for their altered positioning and migration after disruption of the selenoprotein gene. Besides, it was previously reported that SELENOT is required for glycosylphosphatidylinositol (GPI)-anchored protein synthesis (Liu et al., 2018). GPI anchoring of proteins is a post-translational modification that occurs in the endoplasmic reticulum (Ferguson et al., 1985) and concerns about 150 cell surface proteins in mammals, including neural adhesive molecules (Karagogeos, 2003). The role of SELENOT in this process could be related to its effect on protein glycosylation (Hamieh et al., 2017) since N-glycan moieties of GPI-anchored proteins play an important role in the proper inositol deacetylation necessary for their transport and their calnexin/calreticulin quality control cycle. It is thus tempting to speculate that the lack of SELENOT leads to N-glycosylation defaults in GPI-anchored proteins including adhesive proteins present in radial glial fibres and in neuroblasts, which in turn could lead to impairment of glial-cell adhesion and loss of polarity for neuroblasts.

These results obtained after disruption of SELENOT through IUE highlight its crucial role for the proper establishment of projection neuron positioning, through neuroblast polarity maintenance and radial glial scaffolding. Interestingly, analysis of the distribution of profound layer marker CTIP2 shows a mispositioning of deep cortical neurons in Nes-Cre/SELENOT*^fl/fl^* adult mice. These data indicate that the absence of SELENOT impairs the migration of neuroblasts in the neocortex and alters their positioning in the different cortical layers. This finding adds to the list of genes whose expression is essential for the laminar organisation of the neocortex, such as the extracellular matrix protein reelin or the glutamate transporter whose disruption in mice provokes neuroblast stagnation in the ventricular layer, accompanied by a disorganisation of radial glial fibres (Caviness, 1982; Matsugami et al., 2006; Boyle et al., 2011). It is known that reelin binds the apolipoprotein E receptor 2 (ApoER2) in neurons, which also serves as a receptor for SELENOP (a selenium transporter to the brain), thus indicating the possible implication of selenoproteins in important mechanisms involved in cortical lamination since deficiency of ApoER2 in mice leads to abnormal corticogenesis (Hiesberger et al., 1999; Scharpf et al., 2007; Hirota et al., 2018). Besides, absence of adhesion molecules, such as N-glycosylated integrins whose synthesis could be impacted by SELENOT deficiency (Pothion et al., 2020), also leads to cerebral laminar alteration in mice (Anton et al., 1999; Janik et al., 2010). The abnormalities observed in adult SELENOT KO mice on the one hand and after IUE in embryos on the other hand, *i.e* abnormal neuroblast polarity, and alterations of radial glial scaffolding and deep neuron positioning, strongly suggest the importance of SELENOT for the proper laminar organisation of the neocortex by modulating these processes. Since the localisation of neurons in cortical layers is correlated to their identity and connectivities with cortical and subcortical structures (Kwan et al., 2012), it is conceivable that the hyperactive phenotype observed in adult Nes-Cre/SELENOT*^fl/fl^* mice (Castex et al., 2016) could be ascribed to neuron mispositioning induced by alteration of neuroblast migration, but also by exacerbated apoptosis of neuroblasts in the absence of SELENOT, as previously described (Castex et al., 2016).

To conclude, our present data show for the first time that SELENOT is an essential actor in the neocortex required during embryogenesis for cortical lamination, but probably also during the whole mouse lifespan to protect this cerebral structure. We demonstrated the importance of SELENOT during corticogenesis for proper neuroblast migration and positioning. SELENOT could exert its effects by regulating, via redox mechanisms, different cellular processes necessary for proliferation, migration and differentiation of neuroblasts, including folding of adhesive molecules and scaffolding of radial glial cells.

## MATERIALS AND METHODS

### Animals

All animal experiments were approved by the Normandy Ethics Committee on Animal Experimentation (Authorization No. 25444, No. 6637 and No. 16178) and were performed in accordance to the European Committee Council Directives. C57Bl/6J mice were obtained from Janvier Labs (Saint Berthevin, France). SELENOT-floxed (SELENOT*^fl/fl^*) mice were obtained by intern colony breeding of genitors with a SELENOT^fl^ allele produced in collaboration with the Mouse Clinical Institute (Strasbourg, France). Transgenic SELENOT^fl^ mice were generated by homologous recombination of C57Bl/6J genome sequence encoding the active site of the protein, between exons 2 and 3. Nes-Cre/SELENOT*^fl/fl^* KO mice were obtained by breeding of SELENOT*^fl/fl^* mice with Tg (Nestin-Cre)1 Kln +/- knock-in mice, acquired from the Jackson Laboratory (Bar Harbord, ME) and lack of SELENOT in brain of KO mice was previously confirmed (Boukhzar et al., 2016; Castex et al., 2016). Mice were housed under controled conditions with a 12-h light/12-h dark cycle (light on at 07:00 am). Animals had access to food and water *ad libitum* and were maintained at 22 ± 2°C ambient temperature. Embryos were electroporated and/or surgically removed from SELENOT*^fl/fl^* pregnant females, and embryonic stages were determined by the day of appearance of the vaginal plug, as E0.5. Genotypes were obtained by PCR of genomic DNA from tail clips with the following primers: forward, 5’-GGCTTTATGTAAGCAGTTCTAAACTGTTTCTGC-3’; and reverse, 5’-CGCCCCATTTTATAAACTTTGTATGTTTATGCCC-3’, for WT (211 bp) and SELENOT^fl^ (261 bp) alleles; forward, 5’-GGCTTTATGTAAGCAGTTCTAAACTGTTTCTGC-3’; and reverse, 5’-GCCTAGGTTTTACCTGAGAA ACCAAAGG-3’, for SELENOT*^fl/fl^* alleles (409 bp).

### Laser capture microdissection

Mice were anesthetized by 2% isoflurane inhalation and sacrificed by decapitation. Brains were immediately immersed in – 25°C isopentane and stored at – 80°C. For embryos, pregnant females were anesthetized by 2% isoflurane before sacrifice, and embryos head were cryopreserved in tissue freezing medium (Tissue-Tek, Leica) and stored at – 80°C. All steps were done in RNAse-free conditions. Twenty-μm brain coronal sections cut in a cryostat were collected on PEN blades (ZEISS, Göttingen, Germany) and colored with 1% acetate cresyl violet (Sigma-Aldrich, C5042) to discriminate cortical layers, based on the “Atlas of the Developing Mouse brain” (Paxinos) for samples from E12 to post-natal day (PND) 3, and on “Allen Brain Atlas” for samples from PND 7 to 1-year-old. One mm^2^ of each cortical layer was dissected out by the Leica laser capture microdissection apparatus (Leica Microsystems) and collected in a RLT lysis buffer containing 1% β-mercaptoethanol, 4 ng/μL RNA carrier as described by the supplier of the RNeasy® Micro kit (Qiagen, Hilden, Germany).

### Reverse transcription and quantitative real-time PCR

Extraction of RNA was performed using the RNeasy® Micro kit (Qiagen, Hilden, Germany) and the concentration and integrity of RNA were evaluated by picochips (Agilent Technologies, Waldbronn, Germany). Nine ng of total RNA were reverse-transcribed using the QuantiTect® kit (Qiagen, Hilden, Germany). Relative gene expression of SELENOT and NeuN was quantified by real-time PCR (qPCR) with specific primers and the Fast SYBR® Green Master Mix (Applied Biosystems, Courtaboeuf, France). qPCR was carried out on QuantStudio™ Flex’ system (Applied Biosystems), and gene expression was normalized to housekeeping genes (glyceraldehyde-3-phosphate dehydrogenase, GAPDH, and phosphoglycerate Kinase 1, Pgk1) (Boda et al., 2009), by the 2^−ΔΔCt^ method. Primers used were: mouse SELENOT forward, 5’-GGTATCCAGACATCCGCATTG-3’, and reverse, 5’-GGAAAGATGCTATGTGTCTATAAATTGG-3’; mouse NeuN forward, 5’-CTGCCTGTGAGAACCTAGTGCTTT-3’, and reverse, 5’-GGTCCTTCTGATGTCACACTGTTG-3’; mouse GAPDH forward, 5’-CATGGCCTTCCGTGTTCCTA-3’, and reverse, 5’-CCTGCTTCACCACCTTCTTGA-3’; mouse Pgk1 forward, 5’-GTTGCTCCTGAACTCAAATCTCT, and reverse, 5’-GCCCACACAATCCTTCAAG-3’.

### Western blot analysis

Proteins were extracted using a lysis buffer containing 50 mM Tris-HCl, 10 mM EDTA, 1 % Triton and protease inhibitors (Mini tablets Pierce™, ThermoFisher Scientific, Rockford, USA). After centrifugation at 20.000 x g for 15 min, protein content was assayed by the Bio-Rad Protein Assay (München, Germany). Twenty μg of proteins were separated on 10% SDS-polyacrylamide gels and transferred onto polyvinylidene difluoride (PVDF) membranes in a Trans-Blot apparatus (Trans-Blot Turbo™, Bio-Rad, USA). Membranes were then saturated with a buffer containing 5% milk, 0.05% Tween 20 in Tris-buffered saline (TBS) and incubated with the primary antibodies anti-SELENOT (1:1250, Grumolato et al., 2008) and anti-β-Actin (1:5000, ThermoFischer Scientific, MA5-15739) in 5% milk, 0.05% Tween 20 in TBS, overnight at 4°C. The membranes were then incubated with the secondary antibodies goat anti-rabbit peroxidase and goat anti-mouse peroxidase (1:5000, LI-COR, Lincoln, USA) 1 h at room temperature (RT). Visualization of proteins was performed by chemiluminescence (ECL, Bio-Rad, USA) on ChemiDoc (Bio-Rad, XRS+ system).

### RNAscope® *in situ* hybridization

Mice were anesthetized using a solution of ketamine (100 mg/kg) and xylazine (10 mg/kg), and perfused with sterile phosphate-buffered saline (PBS) followed by sterile paraformaldehyde (PFA) 4%. Brains were dissected and post-fixed in PFA 4% overnight at 4°C. A cryoprotection was performed through immersion in sterile sucrose 15% and 30%. Brains were frozen in isopentane in dry ice and stored at – 80°C. Tissues were sectioned into 10-12-μm slices with a cryostat. Slices were washed 5 min in sterile PBS and heated at 60°C in a dedicated oven (ACD HybEZ™ II Hybridization System (220v) with ACD EZ-Batch Slide System, Biotechne, Rennes, France) under Bake mode during 30 min, and then post-fixed during 15 min with sterile PFA 4%. Slices were dehydrated in graded solutions of ethanol at 50%, 70% and twice 100% during 5 min for each bath. Slices were covered with H_2_O_2_ during 5 min and washed with distilled H_2_O two times. Slices were submerged 10 sec in distilled water at 99°C and 5 min in RNAscope 1X Target Retrieval Reagent (ACD Biotechne) at 99°C. Slides were rinsed 15 sec in distilled water at RT, submerged 3 min in ethanol 100% and dried at RT. Slices were covered with protease RNAscope (ACD Biotechne) during 30 min at 40°C and washed with distilled water two times. Hybridization was performed with 50 μl/slice of probe mix during 2 h at 40°C in an oven. Probes used were supplied by ACD Biotechne and were as follows: SELENOT probe targeting the region 1428-3520 (NM_001040396.3), designed in C2; glutamate transporter targeting VGLUT2 (Slc17a6), designed in C3; GABA transporter targeting VGAT (Slc32a1), designed in C3. After hybridization, slices were covered with RNAscope Multiplex FL v2 Amp 1 during 30 min at 40°C and washed with RNAscope wash buffer two times at RT, then covered with RNAscope Multiplex FL v2 Amp 2 during 30 min at 40°C and washed with RNAscope wash buffer two times at RT, and finally with RNAscope Multiplex FL v2 Amp 3 during 15 min at 40°C and washed with RNAscope wash buffer two times at RT (RNAscope Multiplex FL v2 Amp 1,2,3 : RNAscope® Multiplex Fluorescent Reagent Kit v2; ACD Biotechne). After amplification, the staining steps were carried out sequentially. Slices were covered with RNAscope Multiplex FL v2 HRP-C1 (ACD Biotechne) during 15 min at 40°C and washed at RT with RNAscope wash buffer two times, then covered by 80 μL of Opal 520 at 1/1500 (Opal 520 reagent pack, FP1487001KT, AKOYA Biosciences, Malborough, United States) during 30 min at 40°C and washed at RT with RNAscope wash buffer two times, followed by staining of C2 to block with HRP blocker (ACD Biotecne) during 15 min at 40°C and washed at RT with RNAscope wash buffer two times. Slices were covered by 80 μL of Opal 650 at 1/1000 (Opal 650 reagent pack, AKOYA Biosciences) during 30 min at 40°C and washed at RT with RNAscope wash buffer two times, and staining of C3 to block with HRP blocker (ACD Biotecne) during 15 min at 40°C and wash at RT with RNAscope wash buffer two times. Doublecortin was then labelled by immunostaining, as described in the immunohistofluorescence methods. Staining of the nuclei was done with RNAscope® 4-6-diamino-2-phenylindole (DAPI, ACD Biotechne), and the slides were mounted with fluoromount (Sigma-Aldrich). Images were acquired in the Leica Thunder 3D DCV microscope (Leica microsystems) and were processed in ImageJ/Fiji software.

### *In utero* electroporation

*In utero* electroporation was performed in female mice carrying a floxed SELENOT gene (SELENOT*^fl/fl^*), at 14.5 gestational days which corresponds to the peak of neurogenesis (Takahashi et al., 1995). Females were anaesthetised with isoflurane inhalation during surgery. To prevent pain post-injury, buprenorphine at 0.1 mg/kg was injected subcutaneously 10 min before surgery and uterine horn exposition. One μg of control plasmid DNA (pCAG-IRES-eGFP, Megason and McMahon, 2002; Hand et al., 2005) or pCAG-CRE-IRES-eGFP (named pCAG-CRE), where the Cre enzyme was inserted after excision from pSK-Cre1 plasmid (provided by Clifford Tabbin, Harvard Medical school, diluted in PBS containing 0.05% of Fast Green (Sigma-Aldrich, Saint-Quantin Fallavier, France) was injected into the lateral ventricle of embryos using a Hamilton needle (Dutsher, Issy-les-Moulineaux, France). Each embryo was placed between tweezers-type electrodes (CUY 650P1-5, Sonidel, Dublin, Ireland) linked to an electroporator (CUY21 EDIT, Sonidel, Dublin, Ireland) to apply 5 pulses at 50 V/50 ms, separated by 950 ms intervals. Uterine horns were placed back into the mice abdomen and the cavities were sutured-closed.

### Immunohistofluorescence

Embryo heads were collected 24 or 72 h after *in utero* electroporation and fixed in 4% PFA, for 6 h at 4°C. Samples were then dehydrated by immersion in 15% sucrose/PBS overnight at 4°C, and in 30% sucrose/PBS overnight at 4°C before being embedded in tissue-freezing medium (Tissue-Tek, Sigma) and stored at - 80°C. Immunohistofluorescence was performed on 12-μm coronal sections obtained by cryostat microtome (Leica). For SELENOT, nestin and CTIP2 protein staining, slides were post-fixed in 4% PFA for 30 min and blocked in PBS containing 1% bovine serum albumin, 0.3% Triton X-100, 1% donkey serum /for 2 h at RT. The antibodies anti-SELENOT (1:200, Grumolato et al., 2008), anti-nestin (1:250, Neuromics, Edina, USA), anti-DCX (1:250, Santa Cruz Biotechnology, Germany) and anti-CTIP2 (1:100, Invitrogen, Illkirch) were incubated overnight at 4°C. Visualization of immunostaining was performed by Alexa Fluor 650 or 594-conjugated secondary antibodies diluted at 1:500 (Invitrogen). Nuclei were visualized with DAPI (Sigma-Aldrich) at 1 mg/ml in PBS, and slides were mounted with PBS/glycerol 50/50 v/v for observation. Images were acquired with Leica Thunder 3D DCV microscope (Leica microsystems) and were processed using ImageJ/Fiji software.

### Statistical analysis

Data comparisons were performed using the results of at least three independent experiments and were represented as means ± SEM. Statistical analysis was performed on Graphpad Prism 5 software, using Kruskal-Wallis *U* test and Dunn’s post-test, Mann-Whitney *U* test or Unpaired *t* test. Significance was set at \**P*<0.05, \*\**P*<0.01 and \*\*\**P*<0.001.

## Acknowledgments

Images were acquired in PRIMACEN, the cell imaging platform of Normandy.

## Abbreviations

CP: cortical plate
En: embryonic day n
GE: ganglionic eminence
IUE: *in utero* electroporation
IZ: intermediate zone
PND: postnatal day
ROS: reactive oxygen species
SELENOT: selenoprotein T
SP: subplate
SVZ: subventricular zone
VZ: ventricular zone

